# High-Throughput Measurement of Oxidative Stress Indicators: d-ROMs and BAP Assays in a 384-well Plate Format

**DOI:** 10.1101/2025.07.21.665858

**Authors:** Goki Yamada, Sudarma Bogahawaththa, Yuichiro Nishida, Mikako Horita, Kazuhiro Kawamoto, Yasuyuki Maeda, Mikiko Tokiya, Megumi Hara, Tadayuki Tsujita, Akiko Matsumoto

## Abstract

The body’s oxidative balance is regulated by systems that produce and eliminate reactive oxygen species (ROS), and disruptions in this balance can contribute to the development of lifestyle-related diseases. d-ROMs and BAP tests, which evaluate oxidative balance using just 30 µL of serum per sample in approximately 10 minutes, are widely used for this purpose. However, these methods are not suitable for large-scale studies due to their low-throughput and cost. To overcome these limitations, we established a simultaneous measurement system for multiple samples (144 samples) for measuring d-ROMs and BAP using a pipetting robot and a 384-well plate. The developed system demonstrated good linearity and reproducibility, while significantly reducing reagent and sample consumption. Measurement differences compared to the one of the currently available FREE Carrio Duo system ranged from −5% to 3% for d-ROMs and −12% to 8% for BAP, indicating high consistency with existing methods. Furthermore, the measurement time was substantially shortened from 24 hours (10 minutes × 144 samples) to at longest four hours, although the final color measurements for d-ROMs were performed one week later. This optimized semi-automated system enables the precise and efficient measurement of oxidative stress markers, making it suitable for large-scale studies.

## 1. Introduction

Oxidative stress is defined as an imbalance between the production of reactive oxygen species (ROS) and the efficiency of the antioxidative defense system.^1–3^ Increased oxidative stress has been implicated in the development of various diseases, including cardiovascular disease, neurodegenerative disorders, and cancer.^4–6^ The relationship between oxidative stress balance and disease onset is expected to contribute to disease prevention.^7^ Various biomarkers are used to assess ROS production, including thiobarbituric acid reactive substances, serum oxidized low-density lipoprotein, and urinary 8-hydroxy-2’- deoxyguanosine.^8–10^ On the other hand, antioxidative potential is measured by indicators such as superoxide dismutase activity, catalase activity, blood glutathione concentration, and paraoxonase activity.^11,12^ However, detecting these oxidative stress markers typically requires separate samples and specialized testing equipment for each, making it unsuitable for large- scale measurement systems.

Recently, the diacron-reactive oxygen metabolites (d-ROMs) and biological antioxidant potential (BAP) test has gained attention as a method capable of simultaneously measuring both ROS levels and antioxidative potential using the same sample and testing equipment.^13,14^ The d-ROMs test measures hydroperoxides of lipids, proteins, amino acids, nucleic acids, and other biomolecules that have been oxidized by ROS, using a colorimetric reaction.^15^ The BAP test, on the other hand, evaluates the total reducing potential in the body by measuring the ability of antioxidants in the blood to reduce ferric ions (Fe^3^□) to ferrous ions (Fe^2^□), also using a colorimetric reaction.^16^ The d-ROMs/BAP test requires 30 µL of serum per sample and can be performed using a simple procedure that takes approximately 10 minutes.^17^ The latest model, REDOXLIBRA (Wismerll, Tokyo, Japan) can perform the d- ROMs/BAP test simultaneously on a single sample.

However, this equipment is not suitable for large-scale analyses due to the substantial time and manual labor required. Scaling down the d-ROMs and BAP tests is essential for enabling their use in large-scale population studies, clinical trials, and routine health screenings. As oxidative stress and antioxidative capacity are critical biomarkers linked to chronic diseases, aging, and lifestyle factors, there is a growing demand to assess these parameters across thousands of individuals efficiently.^18–20^

To address this limitation, the current study scaled down the d-ROMs and BAP tests and established a high-throughput, semi-automated system incorporating a pipetting robot. This system is capable of simultaneously analyzing 144 samples, enabling the efficient, rapid, and cost-effective measurement of ROS (via d-ROMs) and antioxidative capacity (via BAP). By streamlining the workflow and increasing sample throughput, this advancement significantly improves the feasibility of applying these assays in large-scale studies and routine population-level screenings.

## 2. Materials and methods

### 2.1 Preparation of standard serum for calibration

Whole blood samples were collected from 12 healthy subjects and left at room temperature for 40 minutes. Each serum was obtained through centrifuged at 1,200 × g for 15 minutes and pooled into one tube (approximately 90 mL). Aliquots were stored at −80 °C for future use. d-ROMs and BAP levels were measured using the FREE Carrio Duo (Wismerll, Tokyo, Japan).

### 2.2 Sample preparation and measurement procedure

As shown in Fig. 1, standard serum (STD) was thawed and prepared as a six-concentration dilution series using physiological saline (Otsuka Pharmaceutical, Tokyo, Japan) (2.8- to 13.9-fold dilution) (A). 25 µL of each standard serum dilution was dispensed into a 96-well Semi-Skirted PCR Plate (AZENTA, Burlington, MA, USA: 96-well plate A), as arrangement shown in (B) with yellow backgrounds. 5.0 µL of thawed serum from unknown samples (UKN) were dispensed into in the sample plate (B) with white backgrounds. Then serum was diluted 5-fold by adding 20 µL of physiological saline (C). For the d-ROMs measurement, 10 cuvettes of d-ROMs reagent (Wismerll) were combined with 200 µL of coloring reagent, forming the d-ROMs mixture. For the BAP measurement, 20 cuvettes of BAP reagent were mixed with 1.0 mL of coloring reagent, forming the BAP mixture (D).

**Figure 1.**
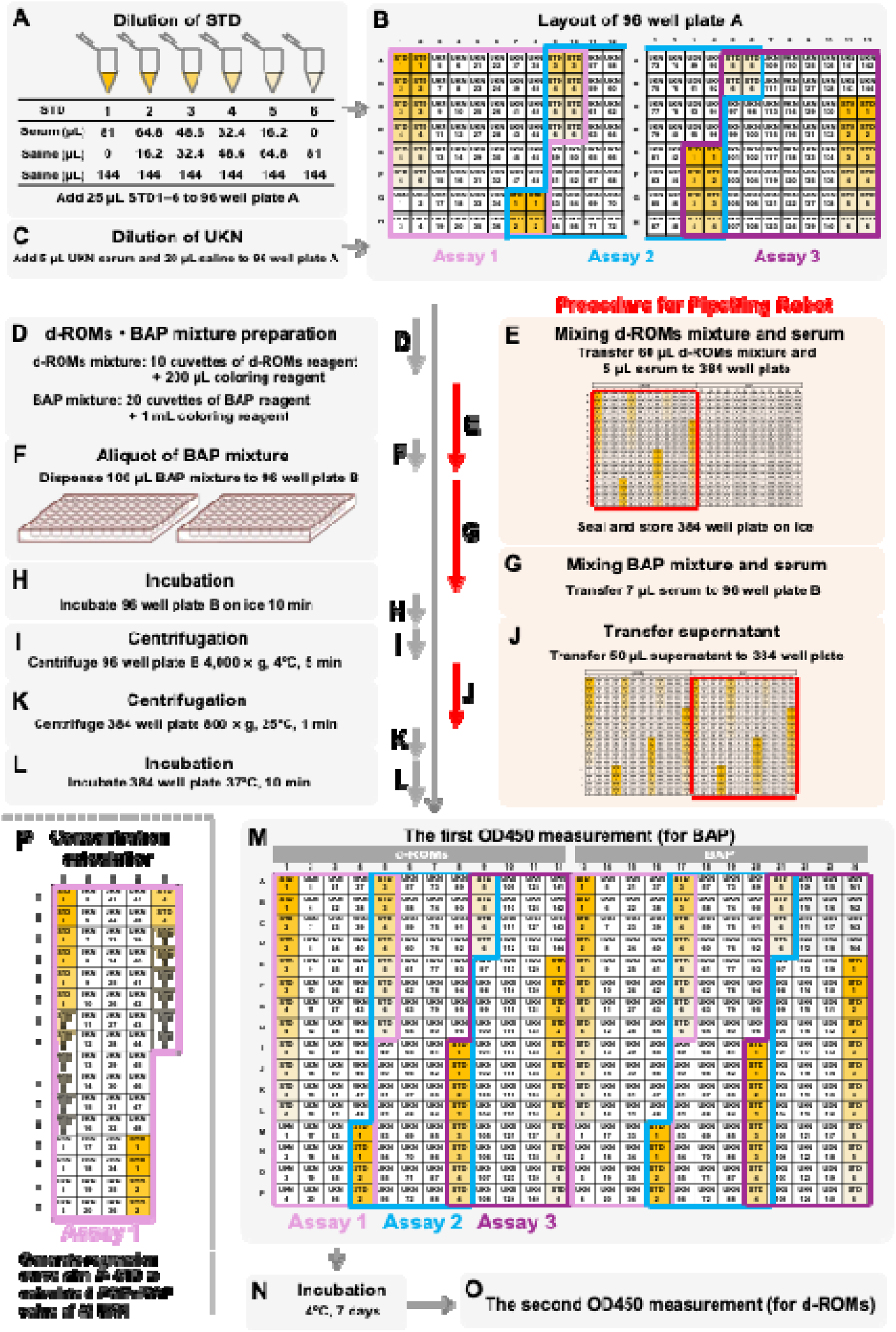
Procedure for measuring d-ROMs and BAP of 144 samples. (A, B) Standard serum (STD) was thawed and prepared as a six-concentration dilution series using saline (2.8- to 13.9-fold dilution). 25 µL of each STD dilution was dispensed into wells with yellow backgrounds in a set of 96-well plates (96-well plate A). (B, C) 5.0 µL of unknown serum (UKN) samples was dispensed into wells with white backgrounds in the 96- well plate A. The UKN was diluted 5-fold by adding 20 µL of saline. (D) For the d-ROMs measurement, 10 cuvettes of d-ROMs reagent were combined with 200 µL of coloring reagent, forming the d-ROMs mixture. For the BAP measurement, 20 cuvettes of BAP reagent were mixed with 1.0 mL of coloring reagent, forming the BAP mixture. (E) 60 µL of the d-ROMs mixture was dispensed into a 384-well plate by a pipetting robot, Andrew+. 5.0 µL of the diluted serum samples from the 96-well plate A was transferred into the 384-well plate by the pipetting robot. After mixing the d-ROMs mixture and serum, the 384-well plate was sealed and temporarily stored on ice. (F) 100 µL of the BAP mixture was manually dispensed into a set of 96-well plates (96-well plate B). (G) 7.0 µL of the diluted serum (remaining samples from the 96-well plate A) were transferred to a 96-well plate B using the pipetting robot. (H) The 96-well plate B was sealed and incubated on ice for 10 minutes. (I) The plate was centrifuged at 4,000 × g at 4°C for five minutes. (J) The seal of the 384-well plate was removed, and 50 µL of the supernatant from the 96-well plate B was transferred to the vacant wells of 384-well plate using the pipetting robot. (K) The 384-well plate was resealed and centrifuged at 800 × g at 25 °C for one minute. (L) The 384-well plate was incubated at 37 °C for 10 minutes. (M) The seal was removed, and the optical density at 450 nm (OD450) was measured by a plate reader for BAP. (N) The plate was then incubated at 4 °C for seven days. (O) The OD450 value was measured again for the d-ROMs. (P) Assay configuration with one STD curve.

A pipetting robot, Andrew+ (Waters, Milford, MA, USA) was used to dispense 60 µL of the d-ROMs mixture into a 384-well plate (PerkinElmer, Waltham, MA, USA). Then, 5.0 µL of the diluted serum samples from the 96-well plate A were transferred into the 384-well plate by the pipetting robot, following the arrangement shown in (E). This resulted in a total volume of 65 µL per well. The mixture was pipetted eight times with 35 µL to ensure thorough mixing. After mixing the d-ROMs mixture and serum using the pipetting robot, the 384-well plate was sealed and temporarily stored on ice. Simultaneously, 100 µL of the BAP mixture was manually dispensed into two 96-well plates (96-well plate B) (F). Then 7.0 µL of the diluted serum (remaining samples from the 96-well plate A) were transferred to a 96-well plate B using the pipetting robot, resulting in a total volume of 107 µL per well (G). The pipetting robot then aspirated and dispensed 60 µL of each sample eight times for proper mixing. Finally, the 96-well plate B was sealed and incubated on ice for 10 minutes (H). The plate was then centrifuged at 4,000 × g at 4°C for five minutes (I).

### 2.3 Absorbance measurement

The seal of the 384-well plate was then removed, and 50 µL of the supernatant from the 96- well plate B was transferred to the vacant wells of 384-well plate using the pipetting robot, as shown in (J). The 384-well plate was then resealed and centrifuged at 800 × g at 25 °C for one minute (K). After incubation at 37 °C for 10 minutes (L), the seal was removed, and the optical density (OD) at 450 nm (OD_450_) was measured by a plate reader (Multiskan FC, Thermo Fisher Scientific, Waltham, MA, USA) for BAP (M). The plate was then incubated at 4 °C for seven days (N), after that the OD_450_ value was measured again for the d-ROMs (O).

### 2.4 Concentration calculation

As shown in Figure 1M, each 384-well plate contains three double assays (assays 1, 2, and 3) for separate d-ROMs and BAP measurements. As shown in Fig. 1P, a single assay consists of 72 measurements: six levels of duplicate standard solutions (e.g., wells 1A–1L in assay 1 for d-ROMs), 48 unknown samples (e.g., wells 1M–4L in assay 1 for d-ROMs), and an additional series of standards (e.g., wells 4M–4P and 5A–5H in assay 1 for d-ROMs). Consequently, one standard curve is generated using 24 standard samples, and the concentration of each unknown sample is calculated based on this curve.

### 2.5 Reproducibility test

To verify the reproducibility of the current protocol, we performed repeated measurements of normal serum A–D, lipemic serum E–F, and hemolyzed serum G–H. The d-ROMs and BAP measurements were initially performed on fresh samples using FREE Carrio Duo.

Measurements of frozen serum A and B, C and D, and E–H were repeated 132 times, 36 times, and 12 times, respectively, within three assays, using the current method.

Measurements of serum A–D were performed on three discontinuous days. Measurements of serum E–H were performed on a single day.

### 2.6 Sample stability test

The effects of freezing and thawing serum on d-ROMs and BAP values were examined. Whole blood was collected from seven healthy subjects, left at room temperature for 40 minutes, and centrifuged at 1,200 × g for 15 minutes to obtain serum (Fig. 2). The d-ROMs and BAP levels in the seven subjects were measured using the FREE Carrio Duo. Each serum sample was dispensed into five tubes containing 100 µL and five tubes containing 10 µL each. All tubes were frozen and stored at −80°C. Tube sets other than the first set (which underwent a single freeze-thaw cycle) were incubated at 25°C for three hours and then refrozen. This freeze-thaw cycle was repeated as shown in Fig. 2, resulting in serum sample sets that had undergone one to five freeze-thaw cycles.

**Figure 2.**
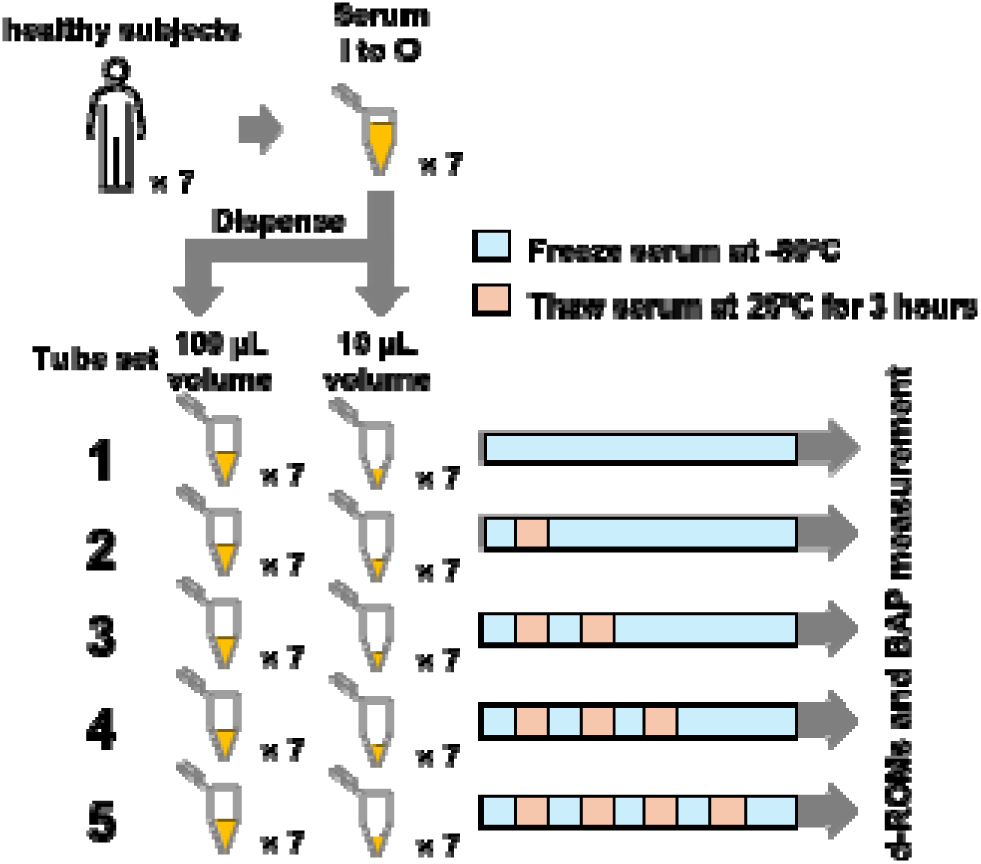
Freezing and thawing of serum for sample stability test. Seven normal sera (Serum I to O) were obtained from seven healthy subjects. Each serum sample was dispensed into five tubes containing 100 µL and five tubes containing 10 µL each. Freeze-thaw cycle was repeated one to five times.

### 2.7 Statistical analysis

Statistical analyses were performed using SAS for Windows version 9.4 (SAS Institute, Cary, NC, USA) and Prism 9 (GraphPad Software, Boston, MA, USA). P values less than 0.05 were considered significant. Dilution linearity and inter-method agreement were assessed using Pearson’s correlation coefficient. The trend effect of freeze-thaw cycle on common-log transformed d-ROMs and BAP was estimated using mixed-effect models that accounted for repeated measurements (proc mixed, SAS 9.4). The Wilcoxon matched-pairs signed rank test was used to detect differences in freeze-thaw cycles or frozen sample volumes.

## 3. Results

### 3.1 Calibration curve

The FREE Carrio Duo analyzer measures OD□□□ however, due to equipment availability, alternative wavelengths were used. As shown in Figures 3Aa and 3Ba, the absorbance spectra of the reaction mixtures were recorded. OD□□□ and OD□□□ were adopted for an available Model 680 microplate reader (Bio-Rad, Hercules, CA, USA) and a Multiskan FC, respectively.

**Figure 3.**
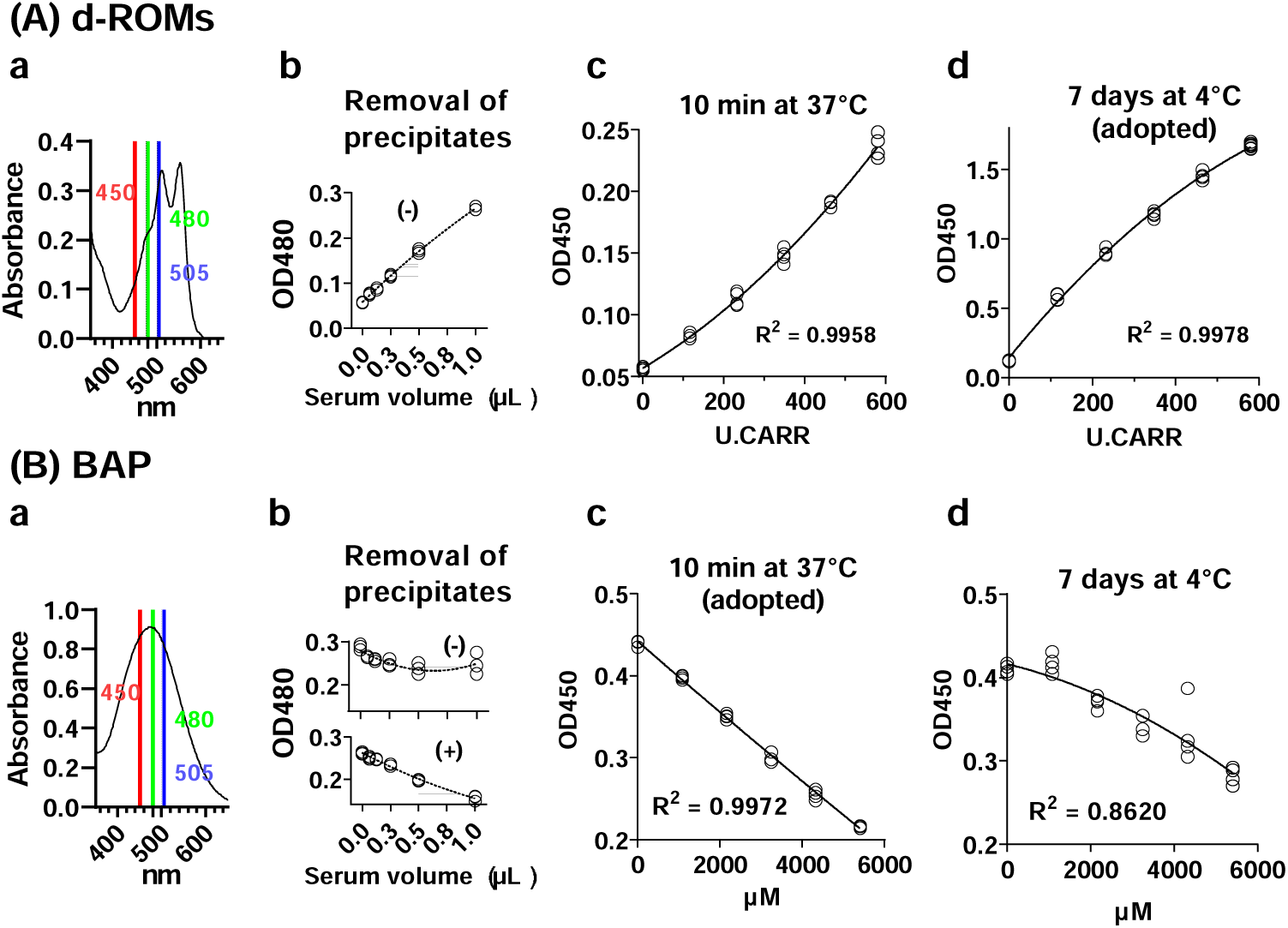
Quantitative determination of d-ROMs (A) and BAP (B) by absorbance. **(a) Absorbance spectra of serum after coloring.** OD_450_ was mainly used in the current study instead of OD_505_, due to the availability of equipment. OD_480_ was used in some preliminary studies. **(b) Dilution linearity at OD_480_.** Absorbance measurements were taken after a 10-minute incubation at 37°C. For BAP, the supernatant was obtained by centrifugation (4,000 × g at 4°C for 5 minutes) to improve linearity. **(c) Dilution linearity at OD_450_.** Measurements were taken after a 10-minute incubation at 37°C. **(d) Dilution linearity at OD_450_.** Absorbance was measured after an initial 10-minute incubation at 37°C, followed by an additional seven-day incubation at 4°C. R-squared values are for second order polynomial.

The formation of precipitates was observed in some sera after mixing with the BAP reagent. As shown in Fig. 3Bb, these precipitates interfered with the dilution linearity of the BAP measurement. To address this issue, the mixture was centrifuged, and the supernatant was used for analysis. This is the absent in FREE Carrio Duo system but significantly improved the dilution linearity. In contrast, d-ROMs measurements showed good dilution linearity even without centrifugation (Fig. 3Ab). Therefore, centrifugation to remove precipitates was necessary only for BAP measurements, not for d-ROMs.

As shown in Fig. 3Ac, the d-ROMs calculation curve after a 10-minute incubation period exhibited good linearity (r² = 0.996). However, better linearity was observed after a seven-day incubation at 4°C (r² = 0.998). Seven days of incubation resulted in higher reproducibility in the high concentration range (Table S1), where d-ROMs have greater diagnostic significance. Based on these observations, a seven-day incubation period at 4°C was adopted for d-ROMs measurements.

For the BAP measurement, after incorporating a centrifugation step and following 10 minutes of incubation at 37°C good dilution linearity was observed (Fig. 3Bc). However, after storing the reaction mixture at 4°C for seven days, the dilution linearity was disturbed (Fig. 3Bd). These results suggest that immediate measurement after incubation is important to ensure accuracy in BAP analysis. Therefore, 10 minutes at 37°C was adopted for BAP measurement.

### 3.2 Reproducibility

The quantitative reproducibility results of serum A–D are summarized in Table 1. The d-ROMs measurements using FREE Carrio Duo ranged from 172 to 486 U.CARR. Subsequently, d-ROMs values were determined using the current method. The intra-assay reproducibility ranged from 1% to 9% coefficient of variation (CV), inter-assay reproducibility ranged from 5% to 9% CV, and inter-day reproducibility ranged from 6% to 8% CV. The differences between the values obtained from the current method and those from the FREE Carrio Duo were within the range of −5% to 3%. These results indicate that the current method provides reproducible measurements, closely comparable to the FREE Carrio Duo system.

**Table 1.** Reproducibility of normal serum.

| Assay number | Day 1 |  |  |  | Day 2 |  |  |  | Day 3 |  |  |  | Day 1-3 |
| --- | --- | --- | --- | --- | --- | --- | --- | --- | --- | --- | --- | --- | --- |
|  | Intra assay |  |  | Inter assay | Intra assay |  |  | Inter assay | Intra assay |  |  | Inter assay | Inter day |
|  | 1 | 2 | 3 | 1-3 | 4 | 5 | 6 | 4-6 | 7 | 8 | 9 | 7-9 | 1-9 |
| A d-ROMs = 204 U.CARR* | (16) | (16) | (16) | (48) | (16) | (16) | (16) | (48) | (12) | (12) | (12) | (36) | (132) |
| Mean | 198 | 208 | 204 | 204 | 214 | 211 | 206 | 210 | 222 | 216 | 210 | 216 | 209 |
| CV | 7% | 5% | 4% | 6% | 7% | 4% | 4% | 5% | 9% | 6% | 4% | 7% | 6% |
| $\Delta^{**}$ | -3% | 2% | 0% | 0% | 5% | 3% | 1% | 3% | 9% | 6% | 3% | 6% | 3% |
| B d-ROMs = 486 U.CARR* | (16) | (16) | (16) | (48) | (16) | (16) | (16) | (48) | (12) | (12) | (12) | (36) | (132) |
| Mean | 457 | 444 | 452 | 451 | 462 | 464 | 476 | 467 | 456 | 471 | 482 | 470 | 462 |
| CV | 7% | 5% | 5% | 5% | 6% | 5% | 5% | 5% | 8% | 6% | 6% | 7% | 6% |
| $\Delta^{**}$ | -6% | -9% | -7% | -7% | -5% | -5% | -2% | -4% | -6% | -3% | -1% | -3% | -5% |
| C d-ROMs = 172 U.CARR* | (4) | (4) | (4) | (12) | (4) | (4) | (4) | (12) | (4) | (4) | (4) | (12) | (36) |
| Mean | 158 | 171 | 159 | 163 | 184 | 169 | 156 | 170 | 176 | 176 | 166 | 173 | 168 |
| CV | 6% | 4% | 4% | 6% | 2% | 1% | 2% | 7% | 2% | 3% | 8% | 9% | 6% |
| $\Delta^{**}$ | -8% | 0% | -7% | -5% | 7% | -1% | -9% | -1% | 3% | 2% | -3% | 1% | -2% |
| D d-ROMs = 176 U.CARR* | (4) | (4) | (4) | (12) | (4) | (4) | (4) | (12) | (4) | (4) | (4) | (12) | (36) |
| Mean | 154 | 181 | 176 | 170 | 168 | 184 | 171 | 175 | 202 | 186 | 171 | 186 | 177 |
| CV | 4% | 2% | 3% | 8% | 4% | 3% | 2% | 5% | 4% | 4% | 1% | 8% | 8% |
| $\Delta^{**}$ | -13% | 3% | 0% | -3% | -4% | 5% | -3% | -1% | 15% | 6% | -3% | 6% | 1% |
| A BAP = 2700 μM* | (16) | (16) | (16) | (48) | (16) | (16) | (16) | (48) | (12) | (12) | (12) | (36) | (132) |
| Mean | 2422 | 2388 | 2421 | 2410 | 2482 | 2478 | 2500 | 2487 | 2507 | 2459 | 2451 | 2472 | 2455 |
| CV | 3% | 3% | 2% | 3% | 3% | 3% | 3% | 3% | 3% | 4% | 4% | 4% | 3% |
| $\Delta^{**}$ | -10% | -12% | -10% | -11% | -8% | -8% | -7% | -8% | -7% | -9% | -9% | -8% | -9% |
| B BAP = 3485 μM* | (16) | (16) | (16) | (48) | (16) | (16) | (16) | (48) | (12) | (12) | (12) | (36) | (132) |
| Mean | 3067 | 3017 | 3161 | 3082 | 3199 | 3218 | 3208 | 3209 | 3281 | 3292 | 3359 | 3311 | 3190 |
| CV | 4% | 7% | 2% | 5% | 2% | 3% | 3% | 3% | 2% | 3% | 5% | 4% | 5% |
| $\Delta^{**}$ | -12% | -13% | -9% | -12% | -8% | -8% | -8% | -8% | -6% | -6% | -4% | -5% | -8% |
| C BAP = 1211 μM* | (4) | (4) | (4) | (12) | (4) | (4) | (4) | (12) | (4) | (4) | (4) | (12) | (36) |
| Mean | 1167 | 1021 | 1015 | 1067 | 1188 | 1052 | 975 | 1072 | 1059 | 1042 | 1115 | 1072 | 1070 |
| CV | 8% | 6% | 4% | 9% | 12% | 3% | 2% | 11% | 3% | 4% | 17% | 11% | 10% |
| $\Delta^{**}$ | -4% | -16% | -16% | -12% | -2% | -13% | -19% | -11% | -13% | -14% | -8% | -11% | -12% |
| D BAP = 1467 μM* | (4) | (4) | (4) | (12) | (4) | (4) | (4) | (12) | (4) | (4) | (4) | (12) | (36) |
| Mean | 1310 | 1316 | 1280 | 1302 | 1355 | 1346 | 1177 | 1293 | 1363 | 1191 | 1386 | 1314 | 1303 |
| CV | 5% | 3% | 7% | 5% | 3% | 6% | 3% | 8% | 6% | 2% | 9% | 9% | 7% |
| $\Delta^{**}$ | -11% | -10% | -13% | -11% | -8% | -8% | -20% | -12% | -7% | -19% | -5% | -10% | -11% |

d-ROMs and BAP of serum A to D were performed for 3 discontinuous days. The number in brackets indicates the number of repetitions. \*d-ROMs or BAP values with FREE Carrio Duo.
\*\* Difference from FREE Carrio Duo.

BAP measurements using the FREE Carrio Duo ranged from 1,211 to 3,485 µM. Intra-assay variation ranged from 2% to 17%, inter-assay reproducibility from 3% to 11%, and inter-day reproducibility from 3% to 10%. The difference between the BAP values obtained using the current method and those measured by the FREE Carrio Duo system ranged from −12% to −8%. These results suggest good reproducibility, with underestimation ranged by 8–12% compared to the FREE Carrio Duo system.

### 3.3 Agreement between FREE Cario Duo and current method

Figure 4 plots the logarithms of d-ROMs and BAP values. The x-axis represents the current method, and the y-axis represents FREE Carrio Duo. The regression equation for log(d-ROMs) is y = 1.05x – 0.11 (R² = 0.9913). For log(BAP), the equation is y = 1.01x (R² = 0.9433).

**Figure 4.**
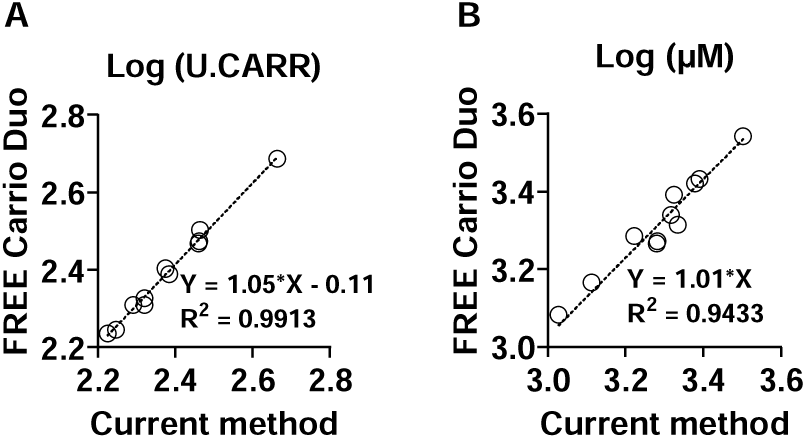
Accuracy of d-ROMs (A) and BAP (B) measurement with the current method. d-ROMs and BAP levels of 11 normal serum A to D and I to O were obtained with FREE Carrio Duo and the current method. R-squared values are for Simple linear regression.

### 3.4 Analyses of lipemic serums

The analysis results for lipemic serum E and F are summarized in Table 2. The d-ROMs values of serums E and F were 378 and 249 U.CARR, respectively, when using FREE Cario Duo. In contrast, the current method with 10 minutes incubation at 37_°C produced significantly higher values, 712 and 536 U.CARR, respectively representing 88% and 115% higher than the FREE Carrio Duo system values. However, with incubation for seven days at 4_°C, the current method yielded values of 419 and 289 U.CARR, respectively, which were much closer to those obtained by the FREE Carrio Duo. These findings highlight the significant impact of lipemia on d-ROMs measurements of the current method. By implementing a seven-day incubation at 4_°C, we were able to minimize this interference and obtain d-ROMs values closely aligned with those measured by the FREE Carrio Duo.

**Table 2.** Reproducibility of lipemic serum.

| Serum E | d-ROMs = 378 U.CARR* | | | | | | | | BAP = 978 μM* | | | |
| --- | --- | --- | --- | --- | --- | --- | --- | --- | --- | --- | --- | --- |
|  | 10 min at 37°C |  |  |  | 7 days at 4°C |  |  |  | 10 min at 37°C |  |  |  |
| Assay number | 1 | 2 | 3 | 1-3 | 4 | 5 | 6 | 4-6 | 1 | 2 | 3 | 1-3 |
|  | (4) | (4) | (4) | (12) | (4) | (4) | (4) | (12) | (4) | (4) | (4) | (12) |
| Mean | 692 | 750 | 693 | 712 | 412 | 433 | 411 | 419 | 1905 | 1756 | 1868 | 1843 |
| CV | 3% | 4% | 6% | 6% | 5% | 4% | 2% | 4% | 8% | 6% | 9% | 8% |
| $\Delta^{**}$ | 83% | 98% | 83% | 88% | 9% | 15% | 9% | 11% | 95% | 80% | 91% | 88% |

| Serum F | d-ROMs = 249 U.CARR* | | | | | | | | BAP = 758 μM* | | | |
| --- | --- | --- | --- | --- | --- | --- | --- | --- | --- | --- | --- | --- |
|  | 10 min at 37°C |  |  |  | 7 days at 4°C |  |  |  | 10 min at 37°C |  |  |  |
| Assay number | 1 | 2 | 3 | 1-3 | 4 | 5 | 6 | 4-6 | 1 | 2 | 3 | 1-3 |
|  | (4) | (4) | (4) | (12) | (4) | (4) | (4) | (12) | (4) | (4) | (4) | (12) |
| Mean | 575 | 474 | 559 | 536 | 286 | 282 | 299 | 289 | 1770 | 1811 | 1670 | 1751 |
| CV | 10% | 4% | 4% | 11% | 1% | 10% | 3% | 6% | 3% | 5% | 6% | 6% |
| $\Delta^{**}$ | 131% | 91% | 125% | 115% | 15% | 13% | 20% | 16% | 134% | 139% | 120% | 131% |

d-ROMs and BAP of lipemic serum E and F were performed. The number in brackets indicates the number of repetitions. \*d-ROMs or BAP values with FREE Carrio Duo. \*\* Difference from FREE Carrio Duo.

According to previous research, normal BAP values for middle-aged and elderly Japanese individuals range from 1,500 to 3,750 µM.^21,22^ In two lipemic samples, as shown in Table 2, the FREE Carrio Duo measured BAP values of 978 and 758 µM, which are notably lower than the expected range. In contrast, the current method yielded values of 1,843 and 1,751 µM, both of which fell within the normal range. These results suggest the robustness of the current method against lipemia, as well as its advantage over the FREE Carrio Duo method.

### 3.5 Analyses of hemolyzed samples

We summarized analytical results for hemolyzed serum G and H in Table 3. The FREE Carrio Duo measured d-ROMs values of 277 and 283 U.CARR, respectively. The current method, after seven days of incubation at 4_°C, yielded similar values of 283 and 290 U.CARR, respectively, showing a 2% difference compared to the FREE Carrio Duo. For BAP measurements, the FREE Carrio Duo yielded values of 2,047 and 2,442 µM, while the current method reported 2,039 and 2,371 µM, respectively; the differences ranged from −3% to 0%. Thus, good agreement was confirmed between the current method and the FREE Carrio Duo for hemolyzed serums.

**Table 3.**
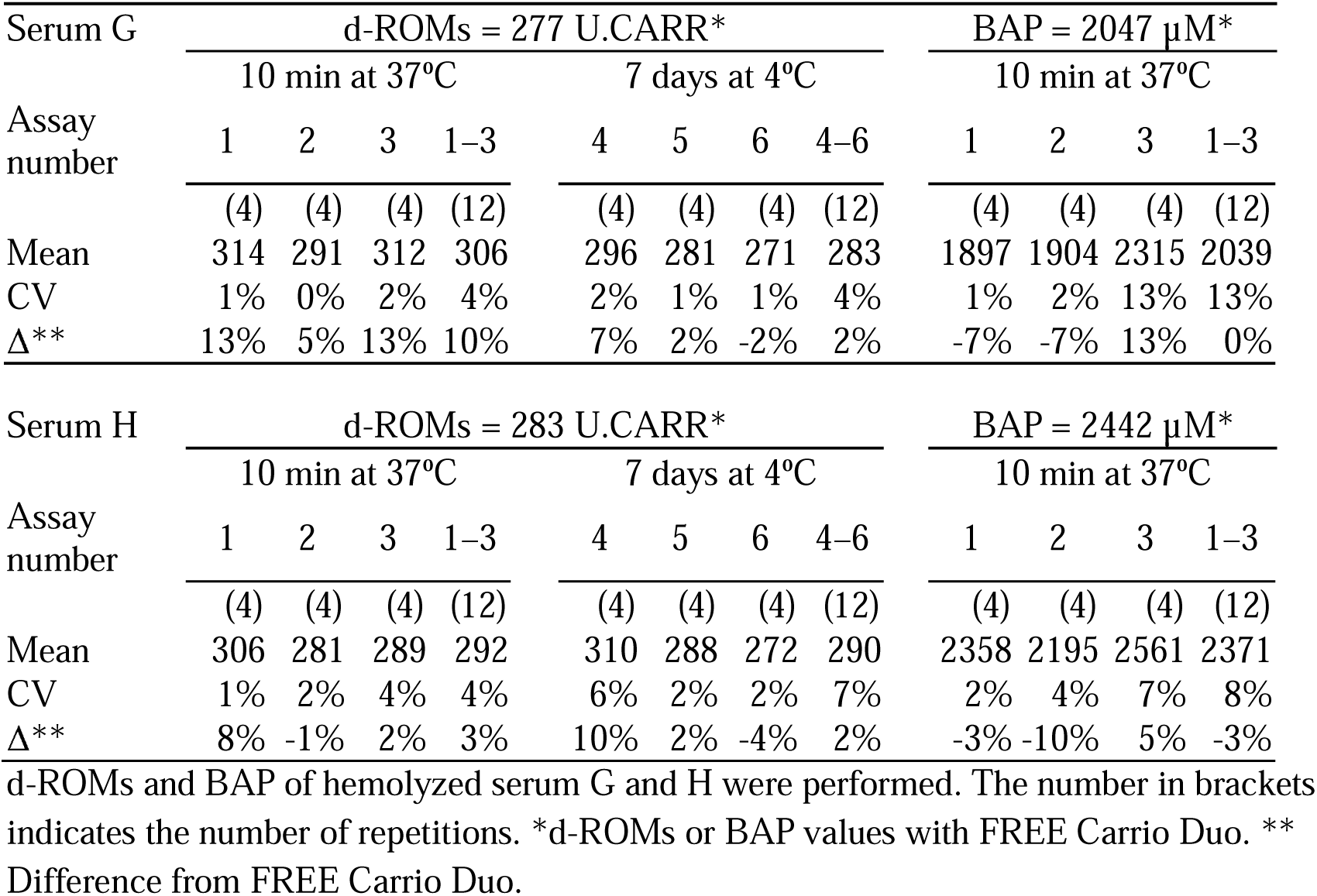
Reproducibility of hemolyzed serum.

| Serum G | d-ROMs = 277 U.CARR* | | | | | | | | BAP = 2047 μM* | | | |
| --- | --- | --- | --- | --- | --- | --- | --- | --- | --- | --- | --- | --- |
|  | 10 min at 37°C |  |  |  | 7 days at 4°C |  |  |  | 10 min at 37°C |  |  |  |
| Assay number | 1 | 2 | 3 | 1-3 | 4 | 5 | 6 | 4-6 | 1 | 2 | 3 | 1-3 |
|  | (4) | (4) | (4) | (12) | (4) | (4) | (4) | (12) | (4) | (4) | (4) | (12) |
| Mean | 314 | 291 | 312 | 306 | 296 | 281 | 271 | 283 | 1897 | 1904 | 2315 | 2039 |
| CV | 1% | 0% | 2% | 4% | 2% | 1% | 1% | 4% | 1% | 2% | 13% | 13% |
| $\Delta^{**}$ | 13% | 5% | 13% | 10% | 7% | 2% | -2% | 2% | -7% | -7% | 13% | 0% |

| Serum H | d-ROMs = 283 U.CARR* | | | | | | | | BAP = 2442 μM* | | | |
| --- | --- | --- | --- | --- | --- | --- | --- | --- | --- | --- | --- | --- |
|  | 10 min at 37°C |  |  |  | 7 days at 4°C |  |  |  | 10 min at 37°C |  |  |  |
| Assay number | 1 | 2 | 3 | 1-3 | 4 | 5 | 6 | 4-6 | 1 | 2 | 3 | 1-3 |
|  | (4) | (4) | (4) | (12) | (4) | (4) | (4) | (12) | (4) | (4) | (4) | (12) |
| Mean | 306 | 281 | 289 | 292 | 310 | 288 | 272 | 290 | 2358 | 2195 | 2561 | 2371 |
| CV | 1% | 2% | 4% | 4% | 6% | 2% | 2% | 7% | 2% | 4% | 7% | 8% |
| $\Delta^{**}$ | 8% | -1% | 2% | 3% | 10% | 2% | -4% | 2% | -3% | -10% | 5% | -3% |

d-ROMs and BAP of hemolyzed serum G and H were performed. The number in brackets indicates the number of repetitions. \*d-ROMs or BAP values with FREE Carrio Duo. \*\* Difference from FREE Carrio Duo.

### 3.6 Effect of sample freezing and thawing

As shown in Fig. 5A, one to five freeze-thaw cycles had no significant effect on d-ROMs values when serum was stored in volumes of either 100 µL or 10 µL. Similarly, BAP values remained stable across repeated freeze-thaw cycles when the serum storage volume was 100 µL. In contrast, when the storage volume was reduced to 10 µL, BAP values increased progressively with the number of freeze-thaw cycles (*P* for trend < 0.001). Samples subjected to four and five freeze-thaw cycles exhibited higher BAP values compared to those thawed once (*P* = 0.054 and 0.013, respectively). As shown in Fig. 5B, d-ROMs values were consistently higher in serum samples stored at a volume of 10 µL compared to those stored at 100 µL across all freeze-thaw cycles. For BAP, no significant difference was observed between the two storage volumes following one to three freeze-thaw cycles. However, after four or five cycles, BAP values were elevated in the 10 µL samples. Based on our observations, the serum storage volume at or above 100 µL is recommended.

**Figure 5.**
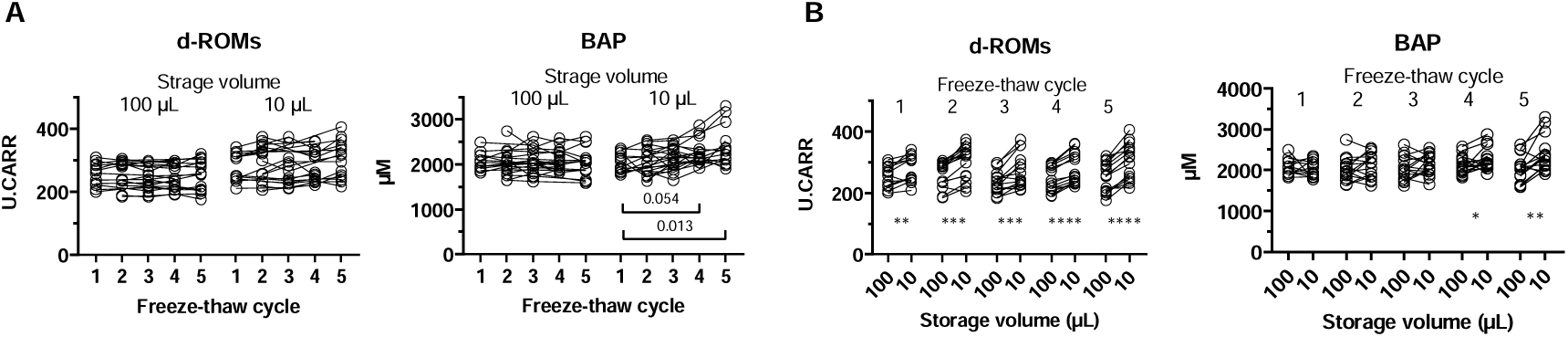
Changes in d-ROMs and BAP values due to freezing and thawing of serum. (A) Variation of d-ROMs and BAP values when 100 µL or 10 µL serum were freeze-thawed 1 to 5 times. P values are for Wilcoxson matched-pairs signed-rank test. (B) Comparisons of d-ROMs and BAP values between serum of 100 µL and 10 µL storage volume at each freeze- thaw cycle. * p < 0.05, ** p < 0.01, *** p < 0.001, and **** p < 0.0001 by mixed-effects models.

## 4. Discussion

In this study, we developed a high-throughput semi-automated system for the simultaneous measurement of d-ROMs and BAP, key indicators for evaluating oxidative balance. By integrating a pipetting robot with a 384-well microplate format, the system enables efficient analysis of up to 144 samples in a single run. The assay exhibited good linearity, reproducibility, and agreement with the conventional FREE Carrio Duo analyzer. These results demonstrate the system’s potential as a reliable and scalable alternative for oxidative stress assessment in large sample cohorts.

Compared to the conventional FREE Carrio Duo, the newly developed method showed several advantages. This new system greatly reduced the amount of sample and reagents needed. The volume of sample required was reduced by 83%, while the d-ROMs and BAP reagents were reduced by 93% and 86%, respectively. As a result, this method saves precious biological samples and lowers overall testing costs. In addition, the total measurement time was significantly reduced from 24 hours (when processing 144 samples individually using the FREE Carrio Duo) to just a few hours with the semi-automated system. Although the final color measurement for d-ROMs is taken after one week, the overall workflow was still much faster. Using a pipetting robot allowed for efficient large-scale testing, even when human resources were limited. This makes the method especially useful for high-throughput studies.

Furthermore, our current method provides a practical solution for testing lipemic samples. Lipemic samples are known to interfere with colorimetric assays,^23,24^ potentially leading to inaccurate measurements. In clinical settings, serum samples often appear cloudy due to elevated postprandial triglycerides levels,^25^ which can affect the accuracy of test results. For this reason, conventional methods advise users to avoid postprandial sampling in order to prevent underestimation of BAP values. For example, the BAP values obtained by FREE Carrio Duo for lipemic samples were 758 and 978 µM, which is far lower than the expected range of 1,500 to 3,750 µM.^21,22^ However, our new method includes a centrifugation step to clarify the serum and produced the BAP values of 1,751 and 1,843 µM, which are within the expected range. Hemolysis is another common issue that can interfere with biochemical assays, including the measurement of d-ROMs and BAP values.^16^ We found no discrepancy between the conventional system and our new systems for hemolyzed sera.

Freeze-thaw cycles are known to induce lipid oxidation and the degradation of endogenous antioxidants.^26^ This can potentially affect the measured levels of d-ROMs and BAP. Additionally, smaller volumes of stored serum are expected to undergo oxidative changes at a faster rate per unit volume, possibly due to increased surface area exposure and reduced buffering capacity.^27^ Our stability tests using 100 µL of serum produced consistent d-ROMs and BAP values, even after five freeze-thaw cycles. However, storing the serum in 10 µL was suggested to be avoided, because BAP values increased with the freeze-thaw cycle, and d-ROMs values were higher than those of the same sample stored in a 100 µL volume. These results suggest that larger serum volumes are more resistant to freeze-thaw degradation and are therefore more suitable for repeated testing.

One limitation of the current method is that the final colorimetric measurement for d-ROMs analysis is performed after one week of incubation. This extended incubation period may not be suitable for applications requiring rapid processing. We have confirmed that for clear serum samples, such as fasting serum, the 10-minute incubation can be applied with reasonable accuracy. However, collecting exclusively fasting serum samples in large cohorts can be challenging. Furthermore, a certain percentage of individuals may have high triglyceride concentrations even in a fasting state, leading to cloudy samples.^28^ Therefore, to broaden the method’s applicability, measurement after a one-week incubation period is recommended. Future research aims to optimize and, if possible, shorten the incubation time to enhance applicability. Another limitation is that we only suggested the method using robotic automation. To further validate and generalize this approach, future studies will compare its performance with manual pipetting.

## 5. Conclusion

In this study, we developed a high-throughput semi automated system for the simultaneous measurement of d-ROMs and BAP using a pipetting robot and 384-well plates. The system was optimized to include lipemic serum samples by incorporating two innovations. First, a centrifugation step was added for BAP analysis. Second, a two-step color measurement was implemented, in which BAP was measured after a 10-minute incubation at 37_°C, followed by d-ROMs measurement after a seven-day incubation at 4_°C. Both d-ROMs and BAP assays demonstrated strong linearity and reproducibility, with minimal variation across repeated measurements. The system demonstrated strong agreement with the conventional FREE Carrio Duo analyzer, with the following regression equations:

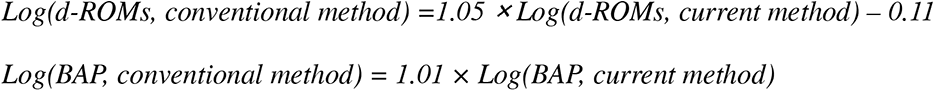

Importantly, the system significantly reduced sample and reagent volumes and shortened measurement time. These improvements make the method both efficient and cost-effective, supporting precise and large-scale analysis of oxidative stress. This system is expected to contribute to the accumulation of valuable data linking oxidative stress with disease, ultimately advancing research in preventive medicine.

## Authors’ contributions

Conceptualization, A.M.; Methodology, A.M.; Investigation, G.Y., S.B., Y.N., M. Horita, K.K., Y.M., and M.T.; Data Curation, G.Y. and S.B; Formal Analysis, G.Y. and A.M.; Writing—Original Draft Preparation, G.Y. and S.B.; Writing—Review and Editing, M. Hara, T.T., and A.M.; Visualization, G.Y. and A.M.; Supervision, Y.N., M. Hara, T.T., and A.M. This research was part of the dissertations submitted by the first author (G.Y.) in partial fulfillment of a Ph.D. degree. All authors have read and agreed to the published version of the manuscript.

## Acknowledgments

We thank Miyuki Fuchigami for her administrative support. We also thank Kanae Mori and Yuka Tokuyama of the Analytical Research Center for Experimental Sciences, Saga University, for technical assistance with analytical equipment.

## Availability of data and materials

The datasets are available from the corresponding author on reasonable request.

## Ethics approval and consent to participate

The study protocol was approved by the Saga University Faculty of Medicine (Project ID: R1-29, R2-31, R3-43, R6-36, R6-42, and R6-53).

---

**Table S1.**
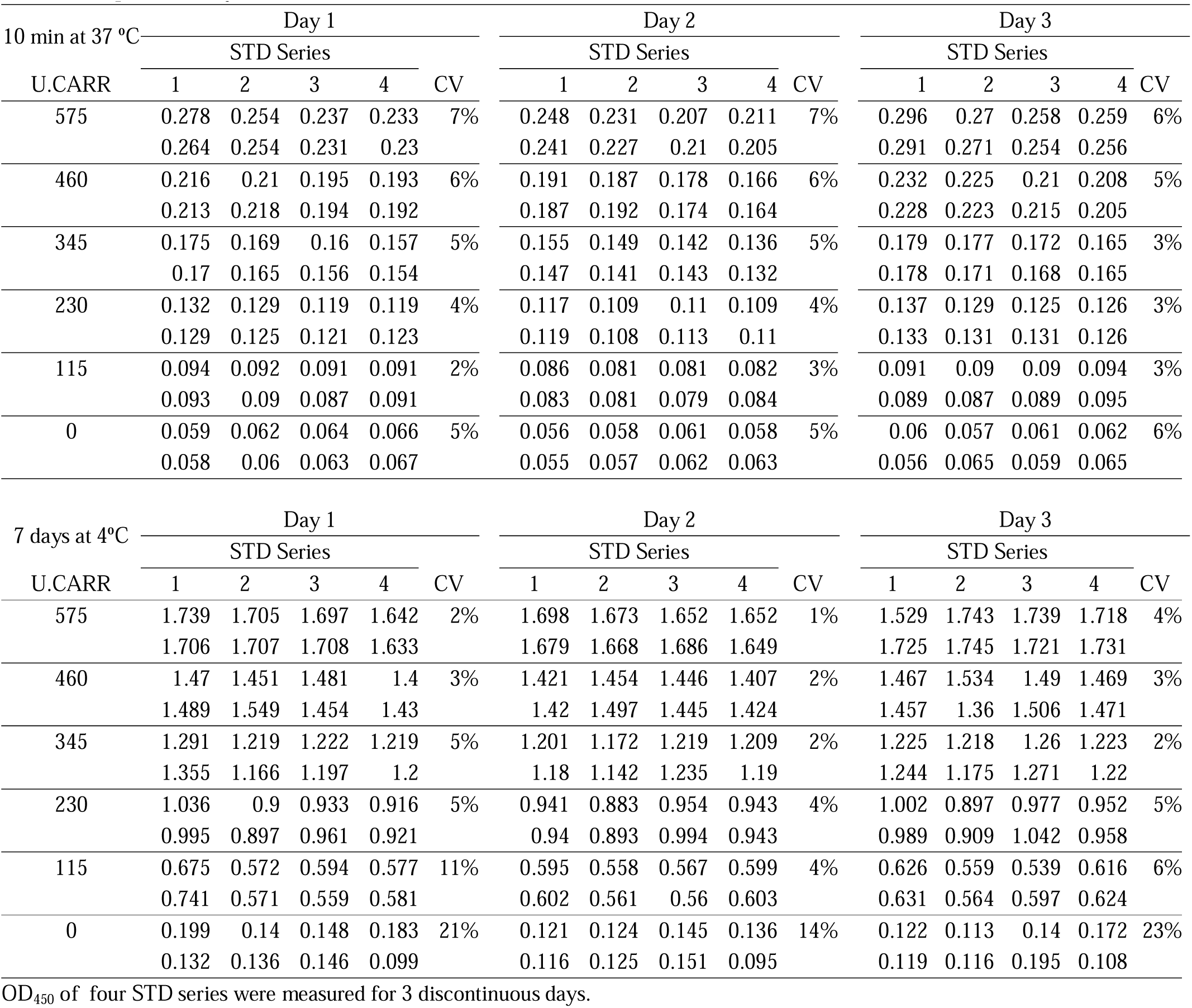
Reproducibility of d-ROMs standards.

